# Pyruvate supports RET-dependent mitochondrial ROS production necessary to control *Mycobacterium avium* infection in human primary macrophages

**DOI:** 10.1101/2022.02.01.478654

**Authors:** Lisa Marie Røst, Claire Louet, Per Bruheim, Trude Helen Flo, Alexandre Gidon

**Affiliations:** Department of Biotechnology and Food Science, Faculty of Natural Sciences, NTNU Norwegian University of Science and Technology, Trondheim, Norway; Centre of Molecular Inflammation Research and Department of Clinical and Molecular Medicine, Faculty of Medicine and Health Sciences, Norwegian University of Science and Technology, Trondheim, Norway

**Author notes:** These authors contributed equally (Lisa Marie Røst, Claire Louet).

**Keywords:** *Mycobacterium avium* infection, innate immunity, human primary macrophages, glycolysis, pyruvate, reverse electron transport, mitochondrial ROS, mitochondrial pyruvate carrier

## Abstract

Macrophages deploy a variety of antimicrobial programs to contain mycobacterial infection. Upon activation, they undergo extensive metabolic reprogramming to meet an increase in energy demand, but also to support immune effector functions such as secretion of cytokines and antimicrobial activities. Here, we report that mitochondrial import of pyruvate is linked to production of mitochondrial ROS and control of *Mycobacterium avium* infection in human primary macrophages. Using chemical inhibition, targeted mass spectrometry and single cell image analysis, we show that macrophages infected with *M. avium* switched to aerobic glycolysis without any major imbalances in the tricarboxylic acid cycle or changes in the energy charge. Instead, we found that pyruvate import contributed to hyperpolarization of mitochondria in infected cells and increased production of mitochondrial reactive oxygen species by the complex I via reverse electron transport, which reduced the macrophage burden of *M. avium*. While mycobacterial infections are extremely difficult to treat and notoriously resistant to antibiotics, this work stresses out that compounds specifically inducing mitochondrial reactive oxygen species could present themself as valuable adjunct treatments.

## INTRODUCTION

Metabolic reprogramming is a key feature of activated macrophages in which the energy production shifts from oxidative phosphorylation to aerobic glycolysis (Beth Kelly 2015). Our understanding of the role of mitochondria recently shifted from sole energy production through oxidative phosphorylation to production of specific signaling metabolites contributing to the anti-microbial and inflammation response. As an example, itaconate, which is a product of the decarboxylation of cis-aconitate by the enzyme Immune Response Gene 1, was placed as a central node that controls immune tolerance and trained immunity ((Jorge Domínguez-Andrés 2019), (Luke A. J. O’Neill 2019)). Further, we and others have independently showed that itaconate, through direct anti-microbial properties, is needed to control viral (Brian P.Daniels 2019), bacterial (Meixin Chen 2020) and mycobacterial infections ((Sharmila Nair 2018), (Alexandre Gidon 2021)). In their seminal paper, Tannahill and colleagues have demonstrated that succinate regulates the production of interleukin 1β via the transcription factor hypoxia induced factor 1α in mouse macrophages challenged with lipopolysaccharide (LPS) (G. M. Tannahill 2013). Later, Mills and colleagues have revealed that mitochondrial reactive oxygen species (mtROS) produced by reverse electron transport (RET) are necessary to initiate and modulate the inflammatory reaction after LPS challenge (Evanna L. Mills 2016). Several papers using undirect approaches also point toward a protective function of glucose metabolism to control mycobacterial infection. Cumming and colleagues reported that *Mycobacterium tuberculosis* (*M. tb*) reduces glycolytic flux to its own benefit (Bridgette M. Cumming 2018); further, Hackett and colleagues proposed that *M. tb* limits glycolytic efficiency by targeting phosphofructokinase activity via the induction of the non-coding RNA miR-21(Emer E. Hackett 2020); and Appelberg and colleagues have suggested that glycolysis is important to control mycobacterial infection in a mouse infection model (Rui Appelberg 2015). Whether and how glycolysis is involved in controlling *M. avium* infection by human primary macrophages remain open questions. Here, we report that macrophages rely on glycolysis and RET to control *M. avium* infection and provide molecular evidence linking pyruvate, the end-product of glycolysis, to anti-mycobacterial mtROS production.

## RESULTS

To examine if primary human macrophages switch to glycolysis after infection with *M. avium* stably expressing DsRed, we first measured glucose consumption and lactate secretion in spent medium by nuclear magnetic resonance. Figure 1 shows a significant increase in glucose uptake (Fig. 1A) and lactate secretion (Fig. 1B), confirming the activation of glycolysis in presence of *M. avium*. Treatment with 100 ng/ml of LPS, a dose reported to shift macrophage metabolism towards glycolysis, increased glucose uptake, albeit not significantly (Fig. 1A, blue), and effectively increased lactate secretion (Fig. 1B). Contrary to Tannahill and colleagues who reported an increased uptake of glutamine during LPS challenge of mouse macrophages (G. M. Tannahill 2013), we did not observe an increase of glutamine consumption in LPS-activated human macrophages (Fig. 1C). The glycolytic switch was further confirmed from targeted mass spectrometry analysis of the intracellular concentration of glucose-6-Phosphate (G6P) and fructose-6-Phosphate (F6P), the first two intermediates of glycolysis, which revealed a significant decrease of both metabolites when compared to the non-infected control (Fig. 1D, white and red). A similar decrease was seen in macrophages treated with LPS (Fig 1D, blue). We next investigated the link between glycolysis and infection by treating infected macrophages with the glycolysis blocker 2-Deoxy-Glucose (2-DG) (Fig. 1E). Images did not show any signs of cell death induced by the treatment. Quantification of *M. avium* intracellular growth after 72 hours of infection revealed a 40% increase in cells treated with 2-DG, showing that glycolysis is indeed required to control the intracellular burden (Fig. 1F).

**Figure 1:**
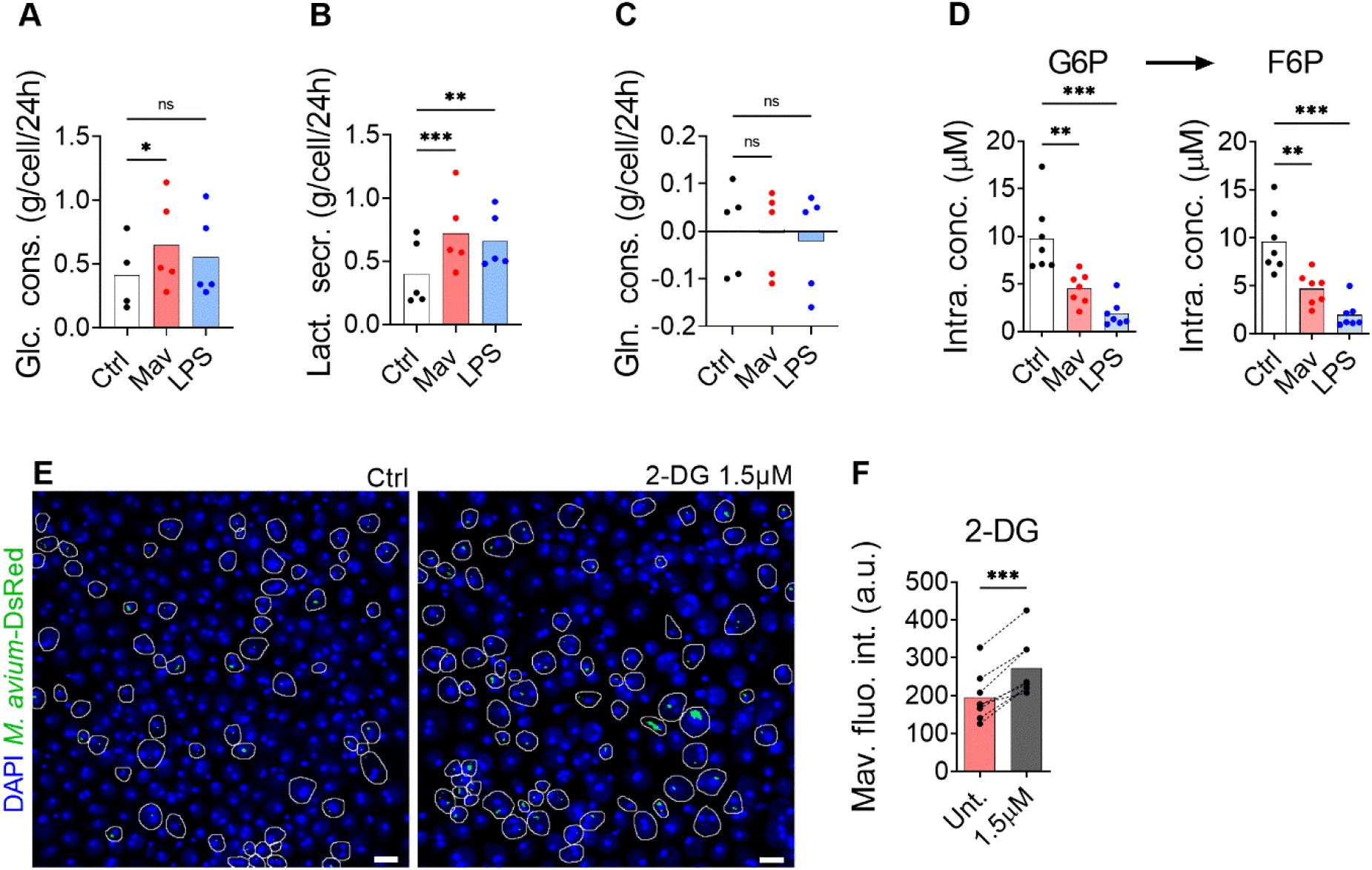
Glycolysis is required to combat *M. avium* infection. Human MDMs were challenged with 100 ng/ml LPS (blue bars) or infected with *M. avium*-DsRed (red bars) for 120 min followed by a chase of 24h (**A-C**). Glucose (Glc) consumption, lactate (Lac) secretion and glutamine (Gln) consumption were measured using nuclear magnetic resonance. Bar-charts represent average from 5 independent donors. Human MDMs were challenged with 100 ng/ml LPS (blue bars) or infected with *M. avium*-DsRed (red bars) for 10 min followed by a chase of 24h (D). Intracellular levels (μM) of glucose-6-phosphate (G6P) and fructose-6-phosphate (F6P) were measured in cell extracts using capillary ion chromatography tandem mass spectrometry. Bar-charts represent the average from 7 independent donors. Human MDMs treated with 2-deoxy-glucose (2-DG) were infected with *M. avium*-DsRed for 10 min followed by a chase of 72h (**E-F**). Intracellular growth was monitored by confocal microscopy. Dots represent the average fluorescence intensity per individual donor (n> 500 cells per donor and per time point), bar-charts represent the average of 7 individual donors. *P* value between untreated and treated conditions were calculated using the non-parametric ANOVA test (**A-D**) or using the non-parametric paired test Wilcoxon signed-rank test (**F**). Scale bars represent 10 μm.

The glycolytic pathway leads to the formation of cytosolic pyruvate. Pyruvate can then be converted into lactate by the lactate dehydrogenase in a reaction simultaneously regenerating NAD^+^ from NADH, into alanine by the alanine transaminase, or it can enter the mitochondria via the Mitochondrial Pyruvate Carrier (MPC) ((Daniel K. Bricker 2012), (Sébastien Herzig 2012)). Mills and colleagues have demonstrated that during LPS activation, mouse macrophages switch to aerobic glycolysis while repurposing the TCA activity to generate specific immunomodulatory metabolites (Evanna L. Mills 2016), which implies that a fraction of the pyruvate formed by glycolysis enter mitochondria. Based on this observation, we hypothesized pyruvate transport into mitochondria is the first step of anti-microbial program centered around mitochondrial metabolism. Quantification by targeted mass spectrometry did not reveal a significant accumulation in the intracellular level of pyruvate in macrophages infected with *M. avium* or treated with LPS when compared to untreated controls (Fig. 2A), suggesting that pyruvate is rapidly metabolized. To investigate the potential role of mitochondrial pyruvate during mycobacterial infection, we treated infected macrophages with the MPC inhibitor UK5099 (John C. W. Hildyard 2005). Quantification of the images revealed a 68% and 46% increase in *M. avium* DsRed signal in cells treated with 100μM and 10μM UK5099, respectively (Fig. 2B), demonstrating that mitochondrial pyruvate is contributing to control *M. avium* infection in macrophages. We have recently demonstrated that IL-6 and TNF-α help controlling the intracellular burden of *M. avium* by supporting the full activation of the Immune Response Factor 1/Immune-responsive gene (IRF1/IRG1) pathway (Alexandre Gidon 2021). To test if the increased burden was not induced by a reduction of TNF-α and IL-6 production, we measured their production upon 2-DG and UK5099 treatment. Blocking glycolysis or mitochondrial import of pyruvate did significantly alter the induction of TNF-α or IL-6 by the infection, demonstrating that pyruvate produced by glycolysis and shuttled in mitochondria contributes to the control of the infection (Fig. 2C and D). Once inside the mitochondrial matrix, pyruvate can be converted into acetyl-CoA by the pyruvate dehydrogenase complex (PDC) (Mulchand S. Patel 2014) and feed into the Tricarboxylic Acid (TCA) cycle producing reducing equivalents for oxidative phosphorylation. Since Ruecker and colleagues have demonstrated that the elevation of fumarate is toxic to *M. tb* (Nadine Ruecker 2017), we decided to probe intracellular concentrations of TCA intermediates. Targeted mass spectrometric quantification showed stable intracellular levels of all measured intermediates after 24h of infection, except for a significant reduction in the level of alpha-ketoglutarate (αKG) (Fig. 2E). Overall, this set of data reveals that no major perturbations of the TCA cycle are induced by the infection, excluding a potential antimicrobial property of these TCA intermediates. We next tested if the infection would affect the energy balance of the cells. Targeted mass spectrometric measurement of the adenine nucleosides AMP, ADP, and ATP, did not reveal any accumulation after 24h of infection, leaving the adenylate energy charge, reflecting the energy status of a cell (D. E. Atkinson 1967), unchanged (Fig. 2F). Since the infection did not alter the energy charge nor the TCA volume, we hypothesized that mitochondrial activity was geared towards mtROS production via the establishment of RET. For this phenomenon to happen, two criteria must be met: the lack of ATP production by oxidative phosphorylation and the maintenance of a high proton motive force/mitochondrial membrane potential ((Evanna L. Mills 2016), (Ellen L. Robb 2018)). To test the engagement of RET, we next treated macrophages infected with *M. avium*-CFP with the potential-insensitive dye MitoTracker Green and the potential-sensitive dye MitoTracker DeepRed ((Enrico Lugli 2005), (Rongbin Zhou 2011), (Andrew W. Greene 2012)) (Fig. 2G). Ratiometric measurement showed that the MitoTracker DeepRed signal was significantly increased by *M. avium* infection and reduced by the MPC inhibitor UK5099 (Fig. 2H), suggesting that pyruvate is involved in the establishment and/or maintenance of high proton motive force induced by the infection. Given that pyruvate import supports the establishment of a high mitochondrial membrane potential and controls *M. avium* infection, we tested whether pyruvate could be necessary to produce mtROS. Measurement of mtROS production using 500nM of the mtROS specific dye MitoSOX (Sergey I. Dikalov 2014) in the presence of the MPC inhibitor (Fig. 3A) revealed a significant reduction of the MitoSOX fluorescence intensity in infected macrophages treated with 10μM UK5099 (Fig. 3B), indicating that pyruvate shuttling into mitochondria is necessary to produce mtROS. Treatment of infected macrophages with various doses of the mitochondrial specific ROS scavenger MitoTEMPO increased the intracellular burden (Fig. 3C), confirming that the production of mtROS is an important factor to control the infection. To confirm that mtROS are produced after the engagement of RET, we exploited the ability of the complex I inhibitor Rotenone to reduce the production of mtROS when RET is engaged (Yulia Kushnareva 2002), (Judy Hirst 2008), (Ru-Zhou Zhao 2019), (Adrian J. Lambert 2004), (Hoi-Shan Wong 2019)). Measurement of mtROS production from macrophages infected with *M.avium*-CFP and co-treated with 10nM of Rotenone revealed a 50% decrease of the MitoSOX signal over the non-treated condition (Fig. 3D), demonstrating both the engagement of RET and the role of complex I during the generation of mtROS. To finally demonstrate the role of RET-related mtROS in the control of the intracellular burden, human macrophages infected with *M. avium* were treated with 10nM Rotenone for 72h. Image quantification revealed a 55% increase of the DsRed signal, indicating a higher intracellular burden (Fig. 3E). The engagement of complex I was confirmed using a high dose of the Carnitine Palmitoyl Transferase 1 inhibitor Etomoxir as an alternative complex I inhibitor (Ajit S. Divakaruni 2018), (Cong-Hui Yao 2018). Treatment of infected macrophage with 100μM Etomoxir increased intracellular burden by 52% after 72h of infection (Fig. 3F) whereas treatment with the complex II inhibitor Dimethyl Malonate (Evanna L. Mills 2016) did not significantly affect the DsRed signal (Fig. 3G). Collectively, these data suggest that pyruvate import into mitochondria contributes to RET-induced mtROS generated by complex I contribute to control the intracellular burden.

**Figure 2:**
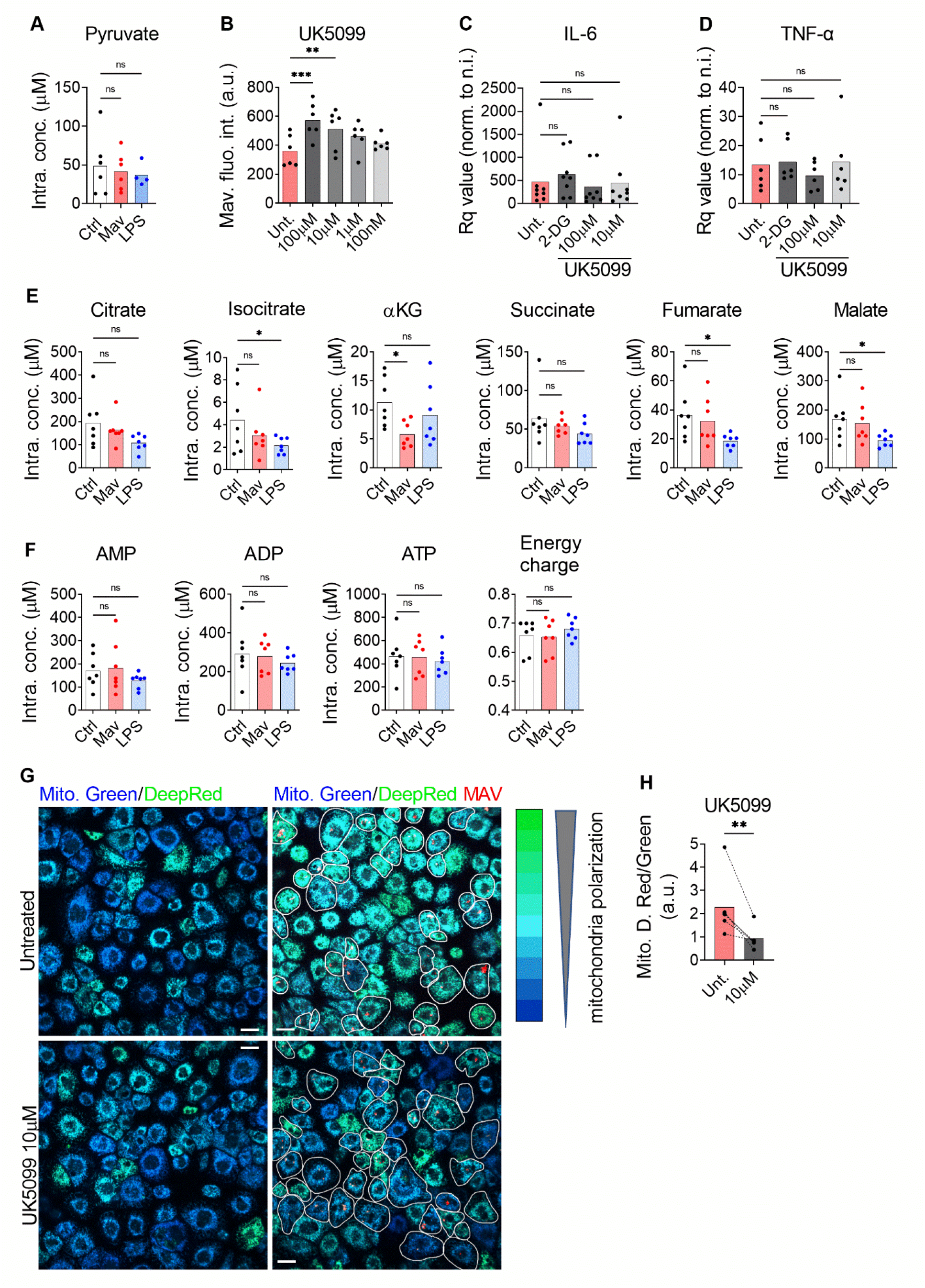
Pyruvate is necessary to maintain mitochondrial hyperpolarization and to control the intracellular burden. Human MDMs were challenged with 100 ng/ml LPS (blue bars) or infected with *M. avium*-DsRed (red bars) for 10 min followed by a chase of 24h (**A**). Intracellular levels (μM) of pyruvate were measured in cell extracts using liquid chromatography tandem mass spectrometry. Bar-charts represent the average from 6 independent donors. Human MDMs were treated with various concentration of UK5099 and infected with *M. avium*-DsRed for 10 min followed by a chase of 72h (**B**). Intracellular growth was monitored by confocal microscopy. Dots represent the average fluorescence intensity per individual donor (n> 500 cells per donor and per condition). Bar-charts represent the average of 6 individual donors. Human MDMs were treated with 1.5 μM 2-DG (grey) or 100 or 10μM UK5099 (grey and light grey) and infected with *M. avium*-DsRed for 10 min followed by a chase of 4h (**C-D**). Induction of IL-6 (C) and TNF-α (D) expression were tested by real-time PCR. Bar-charts represent average Rq values from 8 independent donors. Human MDMs were challenged with 100 ng/ml LPS (blue bars) or infected with *M. avium*-DsRed (red bars) for 10 min followed by a chase of 24h (**E-F**). Intracellular levels (μM) of TCA cycle intermediates (**E**) and adenine nucleosides (**F**) were measured in cell extracts using capillary ion chromatography tandem mass spectrometry. Bar-charts represent average values from 7 individual donors. Human MDMs were treated with 10 μM UK5099 and infected with *M. avium*-DsRed (red) for 10 min followed by a chase of 24h and mitochondria potential was probed using Mitotracker Green (blue) and DeepRed (green) (**G**). Merged images are shown. Infected cells are circled in white. Dots represent the average Mitotracker DeepRed/Green fluorescence intensity ratio per individual donor (n> 250 cells per donor), bar-charts represent the average of 5 individual donors (**H**). *P* value between untreated and treated conditions was calculated using the non-parametric ANAOVA test (**A-F**) and the non-parametric paired test Wilcoxon signed-rank test (**H**). Scale bars represent 10 μm.

**Figure 3:**
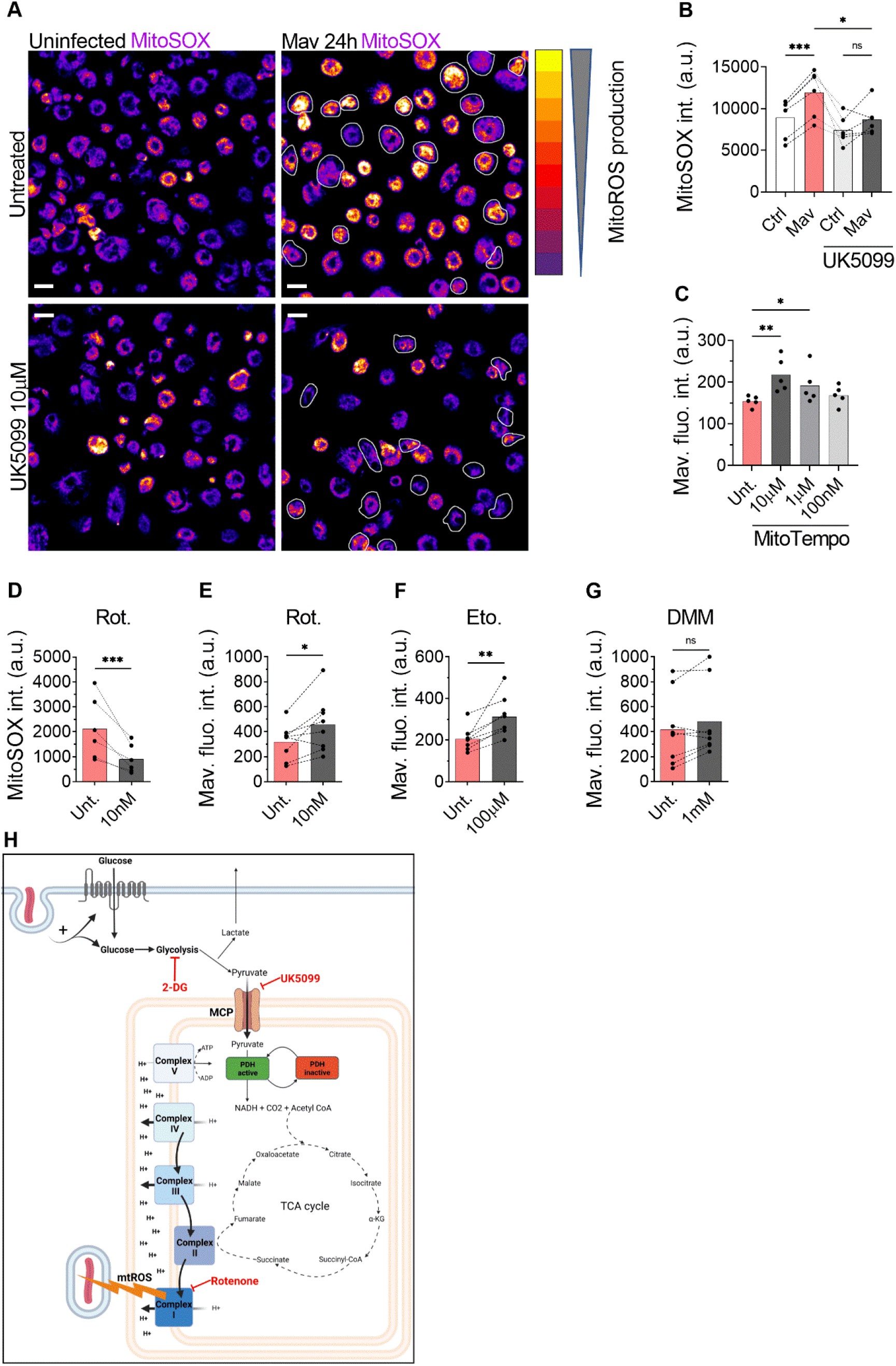
ROS generated by the complex I is necessary to control intracellular burden. Human MDMs were treated with 10 μM UK5099 and infected with *M. avium*-CFP for 10 min followed by a chase of 24h (**A-B**). Mitochondrial ROS were stained using MitoSOX Red. Merged images are shown. Dots represent the average MitoSOX Red fluorescence intensity for uninfected (white bars) and infected (red bars) cells (n> 250 cells per donor). Bar-charts represent the average of 6 individual donors. Infected cells are circled in white. Human MDMs were treated with various concentration of MitoTEMPO (**C**, red). Intracellular growth was monitored by confocal microscopy. Dots represent the average fluorescence intensity per individual donor (n> 500 cells per donor), bar-charts represent the average of 5 individual donors. Human MDMs were treated with 10 nM Rotenone (red) and infected with *M. avium*-CFP for 10 min followed by a chase of 24h (**D**). Mitochondrial ROS were stained using MitoSOX Red. Dots represent the average MitoSOX Red fluorescence intensity (n> 250 cell per donor). Bar-charts represent the average of 6 individual donors. Human MDMs were treated with 10 nM Rotenone, 100 μM Etomoxir or 1mM DMM and infected with *M. avium*-DsRed for 10 min followed by a chase of 24h (**E-G**, respectively). Intracellular growth was monitored by confocal microscopy. Dots represent the average fluorescence intensity per individual donor (n> 500 cells per donor), bar-charts represent the average of 7/8 individual donors. *P* value between untreated (grey) and treated conditions (red) was calculated using the non-parametric ANOVA test (**B-C**) or the non-parametric paired test Wilcoxon signed-rank test (**D-G**). Scale bars represent 10 μm. Working model, generated with Biorender (**H**).

## DISCUSSION

In this manuscript, we confirm that glycolysis is important macrophage defense against mycobacterial infection and describe a central role of pyruvate in linking glycolysis and antimycobacterial mtROS production to control the intracellular burden. Alike Cumming et al. who have demonstrated that the non-pathogenic Bacillus Calmette-Guerin and heat-killed *M. tb* increase glycolysis (Bridgette M. Cumming 2018), we show that human primary macrophages infected with *M. avium* increase glycolysis to facilitate mycobacterial control. We show evidence that the killing mechanism act through the production of mtROS by the complex I of the electron transport chain via the engagement of RET. This mechanism acts in parallel to other immunometabolic defense pathways activated in *M. avium* infected macrophages, such as the production/induction of itaconate via the IRF-IRG1 pathways (Alexandre Gidon 2021). Importantly, whereas we found the IRF1-IRG1 pathway to be regulated by TNF and IL-6, these cytokines did not seem to be involved in regulating the pyruvate-mtROS mechanism described here. Thus, although others have shown that the glycolytic switch in LPS activated macrophages can result in modulation of cytokine responses such as IL-1β, IL-10, our results align with Tannahill and colleagues who also showed that glycolysis inhibition does not alter IL-6 and TNF-α production (G. M. Tannahill 2013) and suggest that macrophages use several mechanisms to control the infection. We show chemical evidence that mitochondrial import of pyruvate through MPC activity is necessary to generate a high membrane potential and the subsequent production mtROS. This finding corroborates with the study of Tai et al, in which the treatment of colon cancer cells with interferon-γ increases MPC1/2 expression and, hence, pyruvate transport into mitochondria to increase mtROS production (YunYan Tai 2019). Unfortunately, our data do not explain how pyruvate is driving RET and mtROS; if pyruvate targets the electron transport chain directly or is converted (via TCA) to another metabolite that initiates RET and mtROS. Several hypotheses have been proposed recently to explain the establishment of RET: 1) In their recent review, Yin and O’neill have proposed that complex II is the site of RET establishment in inflammatory macrophages (Maureen Yin 2021). However, the lack of effect of the complex II inhibitor DMM on *M. avium* burden suggest that complex II is not involved in driving anti-mycobacterial mtROS in our study. 2) It has been proposed that the arrest of complex V drives the electrons to flow backward to complex I and therefore establishes RET (Filippo Scialò 2017). Chin and colleagues have proposed that αKG can bind and inhibit complex V in the model organism *Caenorhabditis elegans* (Randall M. Chin 2014), and αKG was the only TCA metabolite we found significantly changed (decreased) in *M. avium* infected macrophages. However, whether reduced detection of αKG was due to its complexion with complex V and this then, could drive RET and mtROS, remains to be elucidated. Finally, Singhal and colleagues have evaluated retrospectively the effect of Metformin in a cohort of patients infected with *Mtb*. They found that diabetic patients treated with Metformin were less likely to develop severe symptoms triggered by *Mtb* infection.

Using an in vitro model, they demonstrated that, despite reducing the level of glucose, the capacity of Metformin to engage mtROS production was responsible for part of the protective effect (Amit Singhal 2014). With conventional antibiotic-based regimen success rate ranging between 42% and 60% against NTMs infection ((H-B Xu 2014), (Nakwon Kwak 2017), (Roland Diel 2018)), there is an urge to find alternative/combinatory treatment. Our findings suggest a mechanism, summarized in Figure 3H, that links glycolysis and mtROS production during *M. avium* infection in human primary macrophages and bring attention to the possibility of using compounds that specifically engage mtROS production, in adjunct host-directed therapies against mycobacterial infections where lengthy and toxic treatment combined with increasing drug resistance begs for improved therapeutic solutions.

## MATERIAL AND METHODS

### Reagents

The nuclear dye Hoechst 33342 was purchased from Life Technologies. Ultrapure LPS (*E. coli* 0111:B4) was purchased from Invivogen. MitoTEMPO and UK-5099 were purchased from Sigma Aldrich (SML0737; PZ0160). Etomoxir was purchased from Selleckchem (S8244). MitoTracker Green, MitoTracker DeepRed and MitoSOX Red were purchased from ThermoFisher Scientific (M7514; M22426; M36008).

### Isolation and differentiation of human primary macrophages

Buffy coats from healthy blood donors were provided by the Blood Bank, St Olav’s Hospital, Trondheim, after obtaining informed consent and with approval by the Regional Committee for Medical and Health Research Ethics (REC Central, Norway, No. 2009/2245). Peripheral blood mononuclear cells (PBMCs) were isolated using density gradient centrifugation (Lymphoprep, Axis-shield PoC). Monocyte-derived macrophages (MDMs) were generated by plastic adherence for 1h in complete RPMI 1640 (680 μM L-Glutamine and 10 mM Hepes, GIBCO) supplemented with 5% pooled human serum (The Blood Bank, St Olavs hospital) at 37°C and 5% CO_2_. After three washing steps with Hank’s Balanced Salt solution (GIBCO), monocytes were cultivated for 6 days with a change of medium at day 3 in RPMI 1640/10% human serum and 10 ng/ml recombinant M-CSF (R&D Systems). At day 6 the medium was replaced with RPMI 1640/10% human serum and used for experiments on day 7.

### *M. avium* culture, macrophage infection

*M. avium* clone 104 expressing CFP or DsRed was cultured in liquid Middlebrook 7H9 medium (Difco/Becton Dickinson) supplemented with 0.2% glycerol, 0.05% Tween 80 and 10% albumin dextrose catalase. Cultures were maintained at log phase growth (optical density between 0.3 and 0.6 measured at 600 nm, OD600) in a 180-rpm shaking incubator at 37°C for a maximum of 5 days. At the day of infection, bacteria were washed with PBS, sonicated and passed through a Gauge 15 needle to ensure single-cell suspension before challenging day 7 MDMs for 10 minutes at a multiplicity of infection of 10. MDMs were subsequently washed with complete RPMI and maintained in culture for the appropriate time. In some experiments, MDMs were challenged with ultrapure LPS (TLR4; 100 ng/ml) for 24 hours.

### Mitochondria and mitochondria ROS live staining

Human MDMs cultivated on glass-bottomed 96 well plates (IBL) were incubated with 10 nM MitoTracker Green and DeepRed for 10 minutes at 37°C for mitochondria staining or with 500 nM MitoSOX Red for 10 minutes at 37°C for mtROS staining. MDMs were then washed with complete RPMI and immediately imaged with a confocal microscope.

### Imaging

MDMs cultivated on glass-bottomed 96 well plates were imaged with a Zeiss LSM880 confocal microscope with 40x NA=1.4 objective (Carl Zeiss Micro-imaging Inc.). Emissions were collected using GaAsP hybride detectors. The following acquisition parameters were used: 1024*1024 pixel image size, numerical zoom set to 0.6, frame averaging 1, and 3D acquisition to collect the entire cell with a Z-stack step of 0.25 μm. CFP was excited with a 458 nm Argon laser and emissions were collected through a 470-500 nm window. Mitotracker Green was excited with a 488nm Argon laser and emissions were collected through a 505-550 nm window. DsRed and MitoSOX Red were excited with a 543 nm HeNe lasers and emissions were collected through a 560-610 nm window. Mitotracker DeepRed was excited with a 633-diode laser and emissions were collected through a 645-700 nm window. Images were analyzed with Image J (NIH).

### *In situ* CFU measurement, mitochondrial potential and mitochondrial ROS measurements

3D stacks were projected using the “Sum” setting. Resulting images were converted to 8-bit. Regions of interest were drawn around macrophages containing *M. avium*. The background was estimated using HiLo Lookup Tables and subtracted. For mitochondrial potential, Mitotracker DeepRed values were normalized by Mitotracker Green values. A minimum of 250 infected cells per condition and per donor were counted.

### Targeted mass spectrometric quantification of intracellular metabolite levels

Sampling and extraction for mass spectrometric quantification of intracellular metabolites levels was performed as described for adherent cell lines in (Lisa M. Røst 2020). 5-8 million MDMs were sampled for each treatment. Absolute quantification of phosphorylated sugars, TCA cycle intermediates and nucleoside phosphates was performed by capillary ion chromatography tandem mass spectrometry as described in (Hans F. N. Kvitvang 2014) with the modifications described in (Marit H. Stafsnes 2018). Absolute quantification of pyruvate levels was performed by liquid chromatography tandem mass spectrometry with upfront derivatization, as described in (Lisa M. Røst 2020). Data processing was performed in TargetLynx application manager of MassLynx 4.1 (Waters). Absolute quantification was performed by interpolation of calibration curves prepared from serial dilutions of an analytical grade standards calculated by least-squares regression with 1/x weighting. Response factors of the analytical standards and biological extracts were corrected by the corresponding response factor of the U^13^C-labeled isotopologue. Extract concentrations were normalized to seeding density and to the experimental cell volume measured by a Moxi Z automated cell counter to obtain intracellular concentrations.

### Energy charge calculation

Energy charge (EC) was calculated from normalized concentrations of AMP, ADP and ATP using the following formula: EC=(AMP+(0.5*ADP))/(AMP+ADP+ATP) (D. E. Atkinson 1967).

### Quantification of extracellular metabolite levels

Extracellular glucose, lactate and glutamine levels of human MDMs were quantified by recording 1D proton NMR spectra of concentrated fresh and spent (24 hours) medium and applying the TopSpin module ERETIC2 (Topspin 4.1.1, Bruker) as described in (Caroline Krogh Søgaard 2018). The difference between fresh and spent medium was normalized to seeding density to obtain consumption/secretion /cell/24h.

### Statistical analysis

Normality was tested for each experiment. Two-tailed *t*-test and analysis of variance (ANOVA) were used on normally distributed data, Mann and Whitney test was used otherwise. Significant *P* values were set as follows: * <0.05, ** <0.01 and *** <0.005. Statistical analyses were performed using GraphPad Prism 8 (GraphPad Software, Inc., San Diego, CA, USA).

## ACKNOWLEDGEMENTS

This work was supported by grants from the Olav Thon Foundation and the Research Council of Norway (231303, 287696, 223255). All imaging was performed at the Cellular and Molecular Imaging Core Facility at NTNU. Mass spectrometric quantification of central carbon metabolites and nuclear magnetic resonance measurements of extracellular metabolites was performed at the MS and NMR laboratory at the Faculty of Natural Sciences at NTNU, respectively. The summary Figure 3H was created with BioRender.com.

## CONTRIBUTION

AG conceptualized the project. AG and CL did the cell and infection experiments. AG did the microscopy and image analysis with support of CL. LMR did the mass spectrometry. AG, CL, LMR, PB and THF conceived experiments and interpreted results. AG prepared figures. AG wrote the original draft manuscript. All authors reviewed and approved the manuscript.

## Notes

### Competing Interest Statement

The authors have declared no competing interest.

